# *Malassezia* and the Asian menopausal skin

**DOI:** 10.1101/2025.06.24.660996

**Authors:** Bala Davient, Jagpreet Kaur Rikhraj, Aarthi Ravikrishnan, Pritisha Rozario, Cheryl Leong, Ahmad Nazri Mohamed Naim, Nathania Chan, The HELIOS Study Team, John Common, John Campbell Chambers, Yik Weng Yew, Marie Loh, Niranjan Nagarajan, Thomas Larry Dawson

## Abstract

**Background:** Post-menopausal women undergo significant dermatological changes, including thinning skin and reduced sebaceous gland activity, alongside increased incidence of dermatological diseases and hair loss. These changes reshape the skin’s ecological niche, influencing the skin mycobiome composition and behavior. *Malassezia*, a lipid-dependent human pathobiont and dominant fungal resident of skin, has been implicated in several dermatological disorders. We hypothesize that shifts in *Malassezia* populations contribute to post-menopausal skin disorders through altered host–microbe interactions.

**Results:** Shotgun metagenomics of facial and scalp skin from 345 Asian women were stratified by menopausal stage (pre- [N=171], peri- [N=36], and post-menopausal [N=138]) and revealed the presence of seven out of the seventeen recognized *Malassezia* species: *M. globosa*, *M. restricta*, *M. arunalokei*, *M. furfur*, *M. dermatis*, *M. japonica*, and *M. sympodialis*. Detection frequencies of several species varied markedly across menopausal groups. Notably, *M. globosa* was detected 20% more frequently on the scalp of post- versus pre-menopausal women. Reduced sebum concentration on post-menopausal women’s skin correlated with increased *M. globosa* abundance. *In vitro* co-culture of keratinocytes with *Malassezia* spp. showed cells tolerated fungal loads up to 10^4.5^ CFU/cm², but severe cytotoxicity was observed at ≥10^5.5^ CFU/cm². *M. globosa* elicited the highest cytotoxicity towards keratinocytes. All *Malassezia* spp. tested invaded keratinocytes and triggered strong pro-inflammatory responses. Notably, IL-1α, IL-1β, IL-6, IL-8, IL-21, TNF-α, GM-CSF, G-CSF, and MMP1 were significantly overproduced. Transcriptomics of keratinocytes exposed to toxic fungal loads revealed a gene expression profile characteristic of hyperproliferative and undifferentiated cells, alongside elevated expression of *NLRP3*, a key inflammasome sensor involved in pyroptosis.

**Conclusions:** Menopause is associated with distinct shifts in *Malassezia* spp. prevalence and abundance. Reduced skin lipids and thickness may increase fungal burden relative to host cells, promoting inflammation and barrier dysfunction. *Malassezia*’s ability to invade keratinocytes suggests a mechanism for immune evasion and induction of chronic inflammation. Furthermore, keratinocytes exposed to high fungal loads exhibited a transcriptomic profile indicative of hyperproliferation and impaired differentiation, resembling patterns observed in psoriasis, seborrheic dermatitis, and other inflammatory skin conditions. Our co-culture model provides mechanistic insight into *Malassezia*-driven skin inflammation and offers a platform to develop targeted therapies for post-menopausal skin disorders.

## Introduction

Menopause is defined by the permanent cessation of menstrual cycles for at least 12 consecutive months, whether caused naturally, surgically, or chemically. The average age of natural menopause varies by region, with estimates of 47.2 years in Latin America, 47.4 in the Middle East, 48.4 in Africa, 48.8 in Asia, 49.1 in the USA, 50.5 in Europe, and 51.3 in Australia.^1^ As global life expectancy rises and population growth continues, the number of post-menopausal women is expected to increase and remain high.^2,3^ By 2030, approximately half a billion women worldwide will be over 65 years old, a 2.6% increase from 2019.^3^ In Asia alone, this figure is projected to reach approximately 300 million, marking a 3.2% increase from 2019.^3^

During the transition from pre- to peri- and post-menopause, women undergo significant hormonal and physiological changes.^4–6^ Estradiol (E2), the primary estrogen subtype produced by ovarian follicles during reproductive years, declines as follicle reserves diminish. This reduction in serum E2 weakens the hypothalamic negative feedback on gonadotropin-releasing hormone (GnRH) secretion. Consequently, increased GnRH levels overstimulate the anterior pituitary gland, leading to elevated secretion of follicle-stimulating hormone (FSH) and luteinizing hormone (LH) in an attempt to restore E2 levels. However, as ovarian follicles become fully depleted, FSH and LH can no longer stimulate E2 production, resulting in persistently high serum FSH and LH levels alongside reduced E2 levels, marking the hallmarks of menopause.^7,8^ In addition to these hormonal changes, post-menopausal women’s skin becomes thinner, less elastic, less hydrated, and has reduced sebaceous gland activity.^4,5,9,10^ These physiological changes increase the incidence of atopic dermatitis (eczema), psoriasis, and hair loss.^11–14^

Consequently, the physiological changes in post-menopausal women’s skin alter the skin ecological niche and hence the microbial community (the microbiome). Microbial dysbiosis leads to overrepresentation of pathogenic microbes relative to mutualistic/commensal species, increasing susceptibility to skin diseases.^15,16^ While most skin microbiome research in relation to menopause has focused on oral^17,18^ and vaginal^19,20^ epithelia, there is growing interest in other cutaneous regions, such as the face^10,21,22^. Additionally, most skin microbiome studies have centered on bacterial community (bacteriome), leaving menopause-driven changes in the fungal community (mycobiome) largely unexplored. This knowledge gap hinders our understanding of fungal-mediated disorders in post-menopausal women.

The human skin mycobiome is primarily composed of *Malassezia*, a lipid-dependent fungus that thrives in sebaceous gland-rich areas such as the face and scalp.^23,24^ *M. restricta*, *M. arunalokei*, *M. globosa*, *M. furfur*, *M. dermatis*, *M. japonica*, *M. sympodialis*, *M. obtusa*, *M. nana*, *M. yamatoensis*, and *M. slooffiae* are usually found on humans while *M. brasiliensis*, *M. psittaci*, *M. caprae*, *M. equina*, *M. pachydermatis*, *M. cuniculi*, and *M. vespertilionis* are commonly found on animals.^25–28^ Although typically considered a commensal organism, *Malassezia* also has pathogenic potential. In humans, the fungus causes dandruff, seborrheic dermatitis, pityriasis versicolor, and *Malassezia* folliculitis, and is also associated with atopic dermatitis and psoriasis, though its precise mechanistic role remains unclear.^26,29–33^

Post-menopausal women frequently report increased skin sensitivity and itching, which may be influenced by physiological skin changes^11–13^ and potential dysbiosis of the skin mycobiome^34,35^. For example, *Malassezia* secreted aspartyl protease, MfSAP1, degrades human skin extracellular matrix proteins^36^ and may be one of the mechanisms by which *Malassezia* penetrates deeper into the epidermis^37^, thereby leading to skin inflammation. Additionally, *Malassezia* secretes a pseudoprotease, MGL_3331, a putative allergen that elicits increased IgE reactivity in patients with atopic dermatitis.^38^ The fungus also metabolizes polyunsaturated fatty acids present in human sebum, producing various bioactive lipids.^10,39^ Sometimes, the interaction between epidermal cells and *Malassezia* results in the secretion of interaction-specific bioactive lipids. For example, prostaglandin E₂, an inflammatory lipid mediator known to activate mast cells and trigger histamine release^40^, is exclusively secreted by keratinocytes co-cultured with *Malassezia*.^10^

Currently, limited research exists on the prevalence, abundance, and impact of *Malassezia* on women’s skin across different menopausal stages. In this study, we examined a multi-ethnic, Asian-centric female cohort from Singapore (HELIOS)^41^ and found that *Malassezia* dominates the skin mycobiome. Our analysis revealed differences in *Malassezia* prevalence between the cheek and scalp across pre-, peri-, and post-menopausal women. Additionally, we identified species-specific correlations between *Malassezia* and changes in skin physiology. To further validate these associations, we utilized an *in vitro* co-culture model with skin-relevant *Malassezia* species and keratinocytes and showed that cytotoxicity arose beyond a specific fungal dose, with keratinocyte invasion and induction of hyperproliferative, undifferentiated phenotype. The co-culture model offers insights into *Malassezia*-keratinocyte interactions and potential contributions to increased skin sensitivity, itching, incidence of atopic dermatitis, psoriasis, and hair loss commonly reported by post-menopausal women.

## Results

### Serum estradiol and follicle-stimulating hormone classify menopausal stage and highlight differences in skin physiology

We recruited 382 female subjects aged 30 and above in Singapore as part of the Health for Life in Singapore (HELIOS) study and the Asian Skin Microbiome Program (ASMP).^10,41^ After excluding 37 subjects identified as potential confounders in our hypothesis testing (see materials and methods), 345 representative subjects remained (**Fig. 1A**). We hypothesized that serum E2, FSH, and luteinizing hormone (LH), as well as skin sebum, pH, and transepidermal water loss (TEWL), or combination thereof, can replace age as an objective and accurate menopausal stage classifier. Principal component analysis (PCA) of serum E2, FSH, LH, skin sebum, pH, TEWL, and body mass index (BMI) measures identified two distinct clusters along the primary principal component (PC1) (**Fig. 1B**). Notably, serum E2, FSH, and LH were the most influential measures distinguishing the two clusters on PC1 (**Fig. 1C**). The overlap of FSH and LH on the PCA plot indicates that these variables are correlated and therefore, redundant. This was expected because FSH and LH are secreted by the anterior pituitary gland, stimulated by gonadotropin-releasing hormone.^6^ Therefore, using a combination of serum E2 and FSH levels, we classified menopausal stages as follows: pre-menopause (E2 ≥ 110 pmol/L and FSH ≤ 29 IU/L), peri-menopause (E2 < 110 pmol/L and FSH ≤ 29 IU/L or E2 ≥ 110 pmol/L and FSH > 29 IU/L), and post-menopause (E2 < 110 pmol/L and FSH > 29 IU/L). This classification aligned accurately with self-reported questionnaire data (**Fig. 1B** and **Supplementary Fig. 1**).

**Figure 1.**
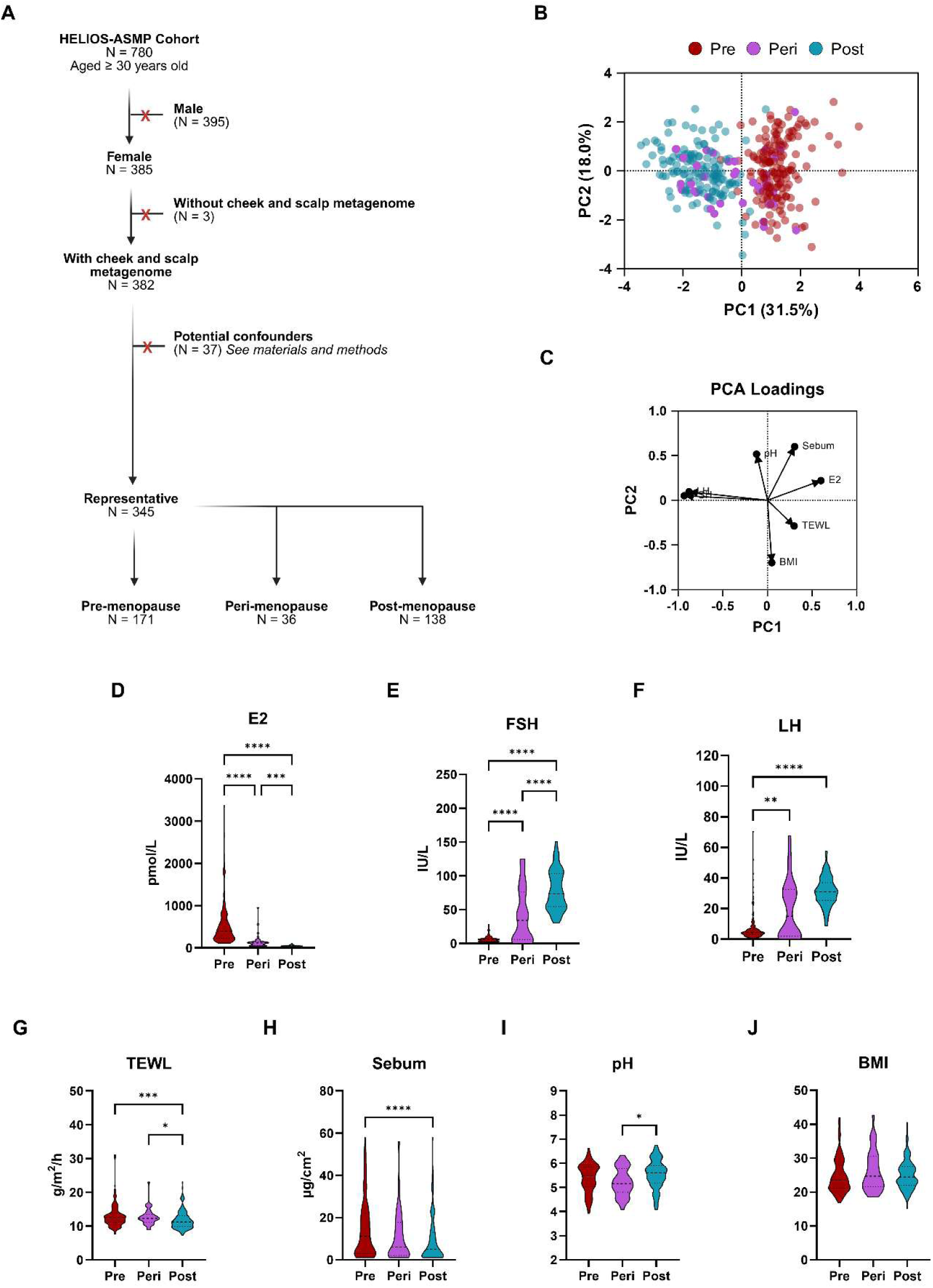
Segmentation between pre-, peri-, and post-menopausal women is driven by differences in female serum hormone concentration and skin physiology measures. (A) Cohort characteristics. Female subjects’ ages ranged between 30 to 75 years old and encompassed Chinese (N = 197), Malay (N = 52), Indian (N = 69), and others (N = 27; Bengali [2], Boyanese [8], Filipino [1], Indonesian [1], Javanese [4], Malayalee [5], Pakistani [2], and Sikh [4]). (B) PCA of representative female serum hormone concentration, skin physiology measures, and BMI. The data was scaled to have a mean of 0 and a standard deviation of 1. PCs were selected based on eigenvalues greater than 1.0 (Kaiser rule). (C) The PCA loadings demonstrate how strongly each variable contributes to the PCs. (D-I) The individual variables used for PCA were re-analyzed using a hormone-based menopause classification approach with serum E2 and FSH levels. The data was analyzed with the Kruskal-Wallis test and post hoc Dunn’s multiple comparisons test. Only significant comparisons are plotted with asterisk. \**p* < 0.5; \*\**p* < 0.01; \*\*\**p* < 0.001; \*\*\*\**p* < 0.0001. See Supplementary **Table 1** for metadata.

This proposed classification of menopausal stages using E2 and FSH identified 171 pre-, 36 peri-, and 138 post-menopausal women among the 345 representative subjects (**Fig. 1A**). As expected, serum E2 concentrations decreased, while FSH and LH levels increased as subjects progressed toward menopause (**Fig. 1D-F**). TEWL, a proxy measure for skin barrier function, was significantly reduced in post-menopausal compared to pre-menopausal women (*p* = 0.0004) (**Fig. 1G**). Additionally, skin surface sebum concentration was lower in post-menopausal compared to pre-menopausal women (*p* < 0.0001) (**Fig. 1H**). Although there was no significant difference in skin surface pH between post-menopausal and pre-menopausal women (*p* = 0.1582), skin pH was significantly higher in post-menopausal compared to peri-menopausal women (*p* = 0.0176) (**Fig. 1I**). This observation was likely due to a transitional decrease in facial skin pH in peri-menopausal women relative to pre- and post-menopausal women similarly reported elsewhere.^42^ The BMI did not significantly differ between groups (**Fig. 1J**). These findings demonstrate an association between the menopausal stage and alterations in skin physiology.

### Malassezia associations with hormone levels and skin physiology

While the composition of the skin bacteriome is known to change significantly as women progress toward menopause,^15,43^ the impact on the mycobiome (the fungal community) remains poorly understood. Building on our previous work, we incorporated updated *Malassezia* genomes (**Fig. 2A**) into a custom fungal database (**Supplementary Table 2**), increased the number of female participants in our cohort, and reanalyzed the sequencing reads from the cheek and scalp of the 345 representative female subjects.

**Figure 2.**
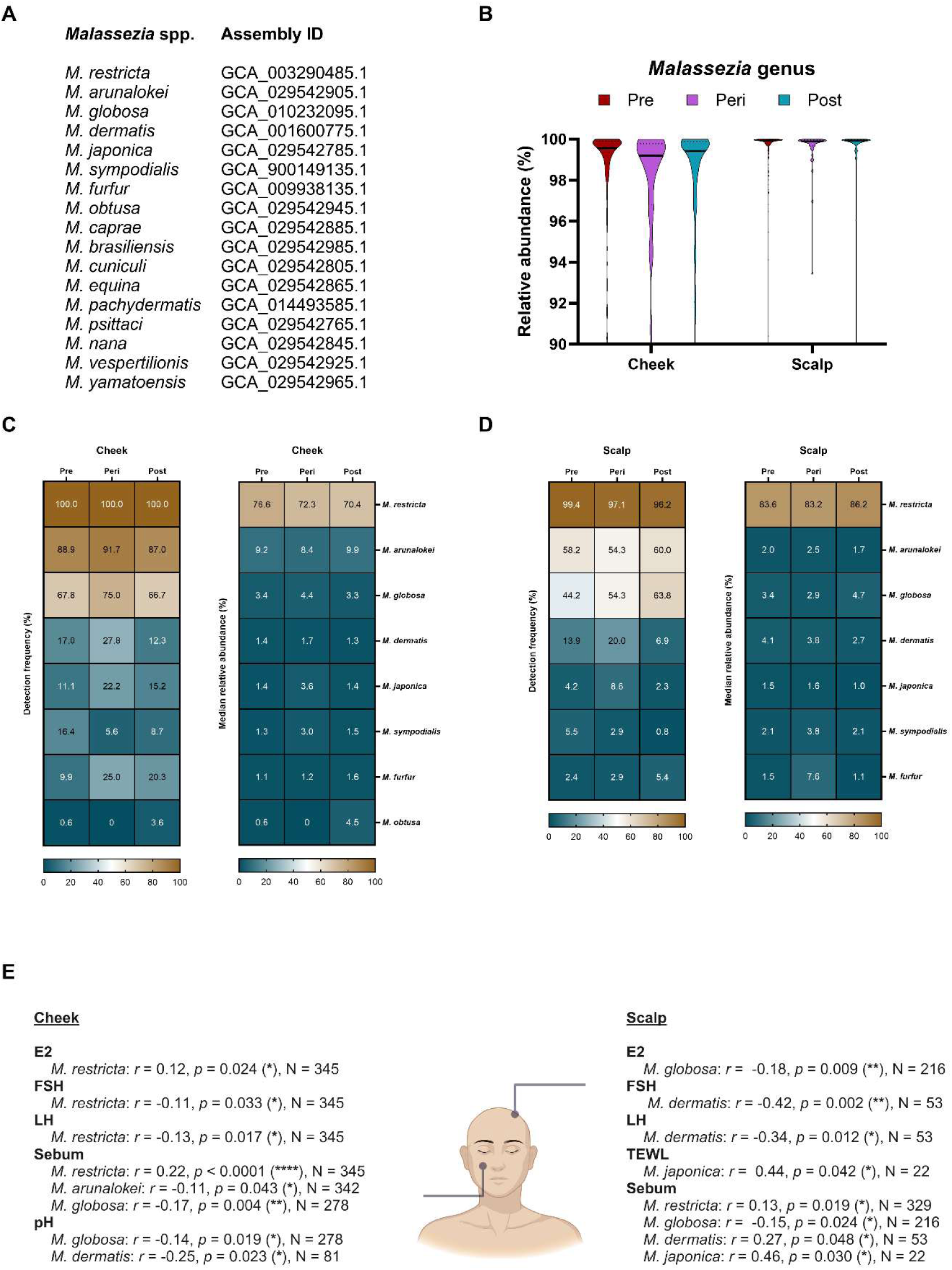
Detection frequency, relative abundance, and correlation of *Malassezia* species with female serum hormones and skin parameters across pre-, peri-, and post-menopausal groups are species-specific. (**A**) Genome assemblies of *Malassezia* species used for shotgun metagenomics analyses. (**B**) Median relative abundance of the *Malassezia* genus relative to the analyzed mycobiome. A two-way ANOVA with post hoc Tukey’s multiple comparison test was performed. Detection frequency (left) and median relative abundance (right) of *Malassezia* species on the (**C**) cheek and (**D**) scalp. See **Supplementary Table 3** for post-menopause versus pre-menopause two-way Fisher’s Exact test. A species was considered as detected if it had a relative abundance of more than 1%. Multiple Mann- Whitney tests with a two-stage step-up (Benjamini, Krieger, and Yekutieli) false discovery rate (FDR) of 5% were used to determine significant differences in relative abundance between pre- and post- menopausal groups for (**C**) and (**D**). (**E**) Correlations between serum hormone levels and skin physiology measures with *Malassezia* species. Correlation analyses were conducted using the Spearman correlation test. \**p* < 0.5; \*\**p* < 0.01; \*\*\**p* < 0.001; \*\*\*\**p* < 0.0001. See **Supplementary Table 2** for the custom fungal genome database used and **Supplementary Table 4** for detailed relative abundances.

The median relative abundance of *Malassezia* at the genus and species levels did not significantly differ across menopausal stages (**Fig. 2B-D**). However, the detection frequency of *Malassezia* species varied notably between pre- and post-menopausal women on both the cheek and scalp. On the cheek, the detection frequency of *M. furfur* increased by 10.4% in post-menopausal women versus pre-menopausal women (**Fig. 2C**). The odds ratio indicates that the detection of *M. furfur* is at least twice as likely on post-menopausal women’s cheeks versus pre-menopausal (Fisher’s exact test odds ratio [FETOR] = 2.3; *p* = 0.0083) (**Supplementary Table 3**). On the scalp, the detection frequency of *M. globosa* increased by 19.6% in post-menopausal women versus pre-menopausal women (**Fig. 2D**). Again, the odds ratio indicates that the detection of *M. globosa* is at least twice as likely on post-menopausal women’s scalp versus pre-menopausal (FETOR = 2.0; *p* = 0.0160). In peri-menopausal women, the detection frequency and relative abundance of *Malassezia* species did not follow a linear trend with menopausal stages, likely due to unstable hormonal levels and progressive changes in skin physiology.

While the median relative abundance of *Malassezia* did not significantly differ between menopausal stages, individual relative abundance values, when paired with serum hormone levels and skin physiology measures, revealed significant associations (**Fig. 2E**). On the cheek, *M. dermatis* was negatively correlated with pH, *M. globosa* was negatively correlated with sebum and pH, *M. arunalokei* was negatively correlated with sebum, and *M. restricta* was positively correlated with E2 and sebum but negatively correlated with FSH and LH. On the scalp, *M. dermatis* was positively correlated with sebum but negatively correlated with FSH and LH, *M. japonica* was positively correlated with TEWL and sebum, *M. globosa* was negatively correlated with E2 and sebum, and *M. restricta* was positively correlated with sebum. These findings highlight significant relationships between individual *Malassezia* species, serum hormones, and skin physiology.

### Malassezia is detrimental to keratinocyte viability beyond a critical fungal load

Reduced serum estrogen is associated with epidermal thinning and an increased risk of dermatoses, including skin infections.^44,45^ We hypothesized that epidermal thinning, caused by reduced keratinocyte turnover, alters the skin microbe-to-keratinocyte ratio, thereby increasing the risk of microbe-associated dermatoses. To test this hypothesis, we co-cultured skin-relevant *Malassezia* species (*M. restricta*, *M. arunalokei*, *M. globosa*, and *M. furfur*) with human immortalized keratinocytes (N/TERT-1) at varying cell population ratios relevant to human skin.^46–48^ Keratinocyte viability was reduced in a dose-dependent manner when co-cultured with *Malassezia* at 10^6^ CFU/well (10^5.5^ CFU/cm^2^) and 10^7^ CFU/well (10^6.5^ CFU/cm^2^), but not at 10^5^ CFU/well (10^4.5^ CFU/cm^2^), indicating a toxic and tolerable fungal dose threshold (**Fig. 3A**). Among the species tested, *M. globosa* CBS 7966 was the most virulent, followed by *M. furfur* CBS 14141, *M. restricta* CBS 7877, and *M. aurunalokei* CBS 13387 (**Fig. 3B**). These results suggest that while keratinocytes exhibit a tolerable fungal dose, toxicity arises when the microbe-to-keratinocyte ratio surpasses a critical threshold.

**Figure 3.**
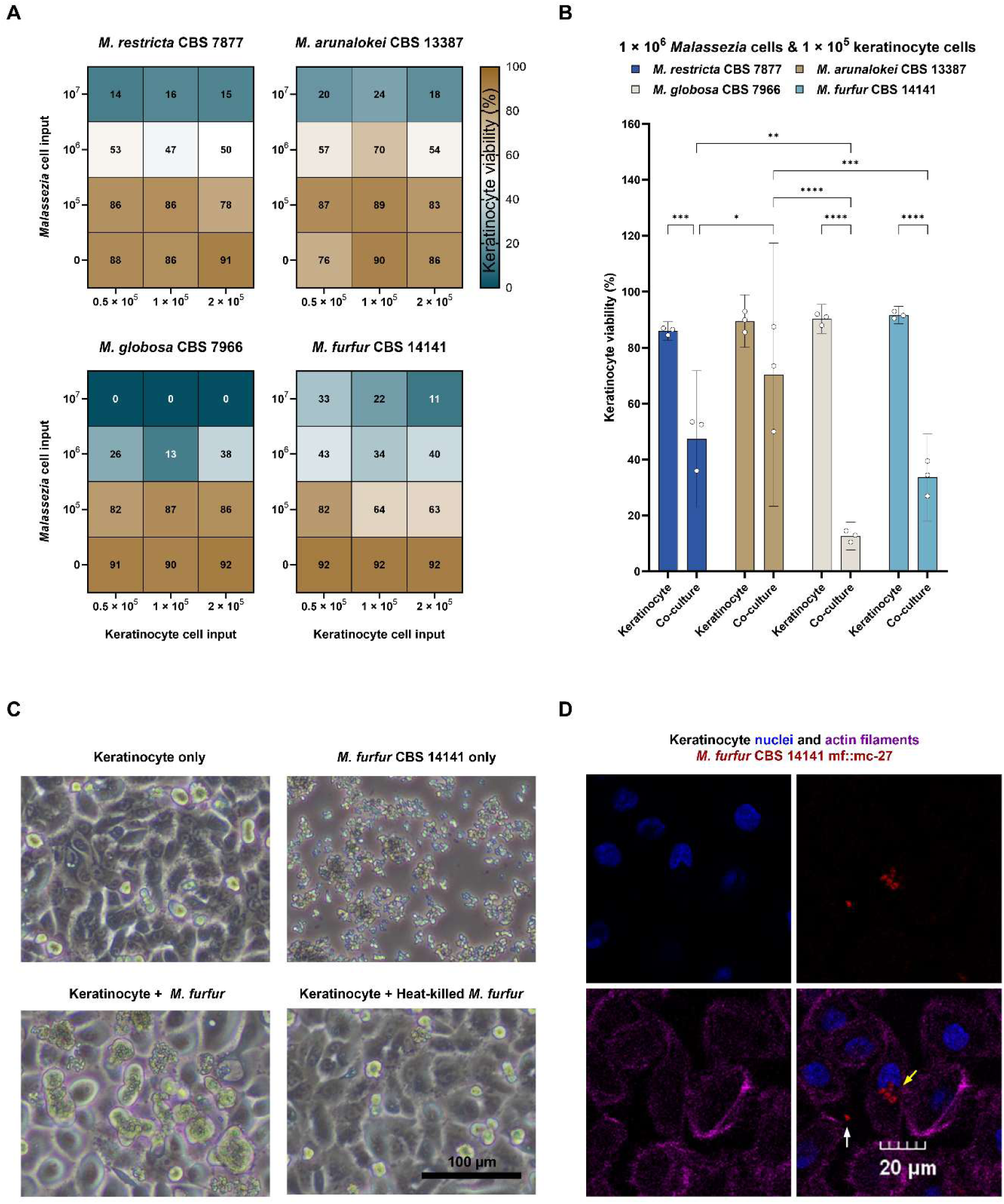
*Malassezia* invades and kills keratinocytes. (**A**) Keratinocyte viability following seven days of mono- or co-culture with *Malassezia* at varying cell population densities. Cultures were maintained at 32°C in a humidified atmosphere with 5% CO₂ in 12-well plates. Keratinocyte viability was assessed using 0.4% trypan blue staining. (**B**) Viability of keratinocytes co-cultured with different *Malassezia* species at a fixed input of 1 × 10^5^ keratinocyte cells/well (10^4.5^ CFU/cm^2^) and 1 × 10⁶ *Malassezia* CFU/well (10^5.5^ CFU/cm^2^). A two-way ANOVA with post hoc Tukey’s multiple comparison test was used (N = 3). \**p* < 0.5; \*\**p* < 0.01; \*\*\**p* < 0.001; \*\*\*\**p* < 0.0001. (**C**) Morphological changes in keratinocytes after seven days of culture with 1 × 10⁶ *Malassezia* CFU/well (10^5.5^ CFU/cm^2^) live or heat-killed mCherry tagged *M. furfur* CBS 14141 (mf::mc-27). Heat-killing was achieved by incubating mf::mc-27 at 60°C for 30 minutes. No CFU were recovered from heat-killed samples, indicating successful killing of *Malassezia* yeast. Differential interference contrast microscopy images are representative of three independent repeats and were acquired at 10× magnification. (**D**) Keratinocytes co-cultured with 1 × 10⁶ *Malassezia* CFU/well (10^5.5^ CFU/cm^2^) live mf::mc-27 for 24 hours. Samples were fixed with 4% paraformaldehyde and stained with DAPI (binds to adenine-thymine-rich DNA regions; pseudocolored blue) and phalloidin (binds to F-actin; pseudocolored purple). mf::mc-27 was pseudocolored red. Yellow and white arrows indicate intra- or extracellular mf::mc-27, respectively. Confocal microscopy images are representative of three independent repeats and were acquired at 40× magnification.

After seven hours of incubation, several keratinocyte cells exhibited an enlarged, globular morphology when co-cultured with live mCherry-tagged *M. furfur* CBS 14141 (mf::mc-27) (**Fig. 3C**). Keratinocyte enlargement was also observed with wildtype *M. furfur* CBS 14141 and other *Malassezia* species tested (data not shown). Confocal laser scanning microscopy revealed that *Malassezia* infiltrated the keratinocytes, localized near the nucleus, and replicated, partially explaining the observed keratinocyte enlargement (**Fig. 3D** and **Supplementary Fig. 2**).

### Malassezia induce a pro-inflammatory keratinocyte response

To assess the inflammatory response of keratinocytes to a toxic dose of *Malassezia* cells, we first exposed keratinocytes to the highest non-lethal fungal dose (10^6^ CFU/well; 10^5.5^ CFU/cm^2^). We then compared cell-free supernatants from co-cultures to those from monocultures using a comprehensive 65-target ELISA-based human immune monitoring panel (**Fig. 4**). In addition, we analyzed the keratinocyte’s transcriptome via RNAseq to determine differentially expressed immune-associated genes (**Supplementary Fig. 3** and **Supplementary Table 5**).

**Figure 4.**
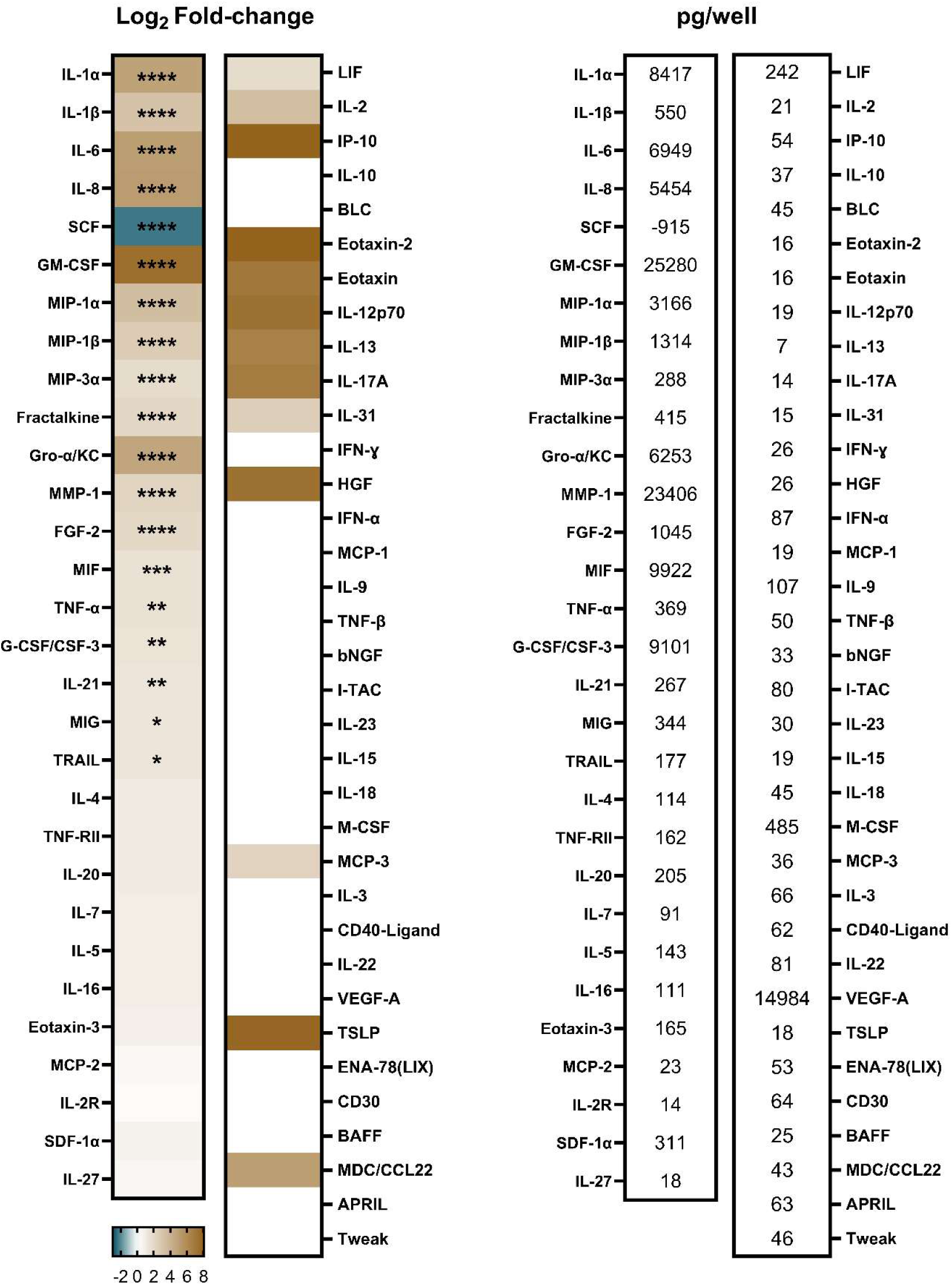
Cytokine profile of keratinocytes co-cultured with *M. furfur* CBS 14141 versus keratinocyte monoculture. Cell-free supernatants from keratinocyte cultures (input: 1 × 10^5^ cells/well; 10^4.5^ cells/cm^2^) incubated for seven days, with or without *Malassezia* (input: 1 × 10^6^ CFU/well; 10^5.5^ CFU/cm^2^), were analyzed using an enzyme-linked immunosorbent assay. Log₂ fold-change values were calculated by dividing the cytokine concentrations of the co-culture by those of the monoculture. A two- way ANOVA with post hoc Bonferroni’s multiple comparison test was performed (N = 3). \**p* < 0.5; \*\**p* < 0.01; \*\*\**p* < 0.001; \*\*\*\**p* < 0.0001. Absolute cytokine concentrations (pg/well) were determined by subtracting the mean monoculture values from the mean co-culture values. Cytokine concentrations for the media-only and *Malassezia*-only conditions were below the detection limit. Brown: high cytokine production; Teal: low cytokine production.

*NLRP3*, an inflammasome encoding gene, was overexpressed over four log-folds in co-cultured keratinocytes versus monoculture, suggesting a high likelihood of pyroptosis, “an inflammatory form of lytic programmed cell death”^49^. Keratinocytes in co-culture also overproduced cytokines IL-1α, IL-1β, IL-6, IL-21, tumor necrosis factor alpha (TNF-α), granulocyte-macrophage colony stimulating factor (GM-CSF or CSF2), granulocyte colony stimulating factor (G-CSF or CSF3), TNF-related apoptosis inducing ligand (TRAIL or TNFSF10), and macrophage migration inhibitory factor (MIF). Furthermore, *IL-23A* gene expression was upregulated in the co-culture.

In terms of chemokines, co-cultured keratinocytes secreted significant amounts of interleukin 8 (IL-8 or CXCL8), melanoma growth stimulating activity alpha protein (Gro- α or C-X-C motif chemokine ligand 1: CXCL1), monokine induced by gamma interferon (MIG or CXCL9), fractalkine (CX3CL1), macrophage inflammatory protein 1 alpha (MIP-1α or CCL3), MIP-1β (or CCL4), and MIP-3α (or CCL20) versus keratinocyte monoculture. Chemokine *CXCL10* gene expression was also upregulated in co-culture versus monoculture, but was not significantly overproduced (IP-10) in the co-culture. In addition, cytokine receptors interleukin 1 receptor type 2 (*IL1R2*) and interleukin 1 receptor like 1 (*IL1RL1*) gene expression were upregulated in co-cultured keratinocytes.

Keratinocytes in co-culture also overproduced proteinases such as matrix metalloproteinase 1 (MMP1) and had upregulated gene expression for *MMP2*, *MMP9*, *MMP10*, and cathepsin S (*CTSS*). While reduced production of stem cell factor (SCF) and increased production of fibroblast growth factor 2 (FGF-2) were noted in co- cultured keratinocytes as part of the 65-target panel, these findings do not provide information regarding inflammation. Overall, the transcript and protein-based analyses revealed that keratinocytes had a pro-inflammatory response to *M*. *furfur* CBS 14141 at a toxic fungal load.

### Keratinocytes exposed to Malassezia show gene expression changes correlated with increased division and decreased differentiation

Pathway analysis of differentially expressed genes predicted activation of cell division pathways in keratinocytes co-cultured with *Malassezia*. Activated pathways include ‘cell cycle checkpoints’ (Z-score: 6.9), ‘mitotic prometaphase’ (Z-score: 6.6), and ‘mitotic metaphase and anaphase’ (Z-score: 6.4) (**Fig. 5**). Some genes were commonly represented in separate pathways. Therefore, we compiled and re- categorized the top 100 up- and downregulated genes within the top 20 significantly perturbed pathways (**Fig. 5**) into simplified biological functions (**Supplementary Fig. 3**). The majority of differentially expressed genes associated with ‘cell division’ were related to DNA replication and mitotic spindle organization. Likewise, downregulation of keratinocyte differentiation genes contributed to the predicted inhibition of the ‘keratinization’ pathway (Z-score: -5.0) (**Fig. 5**). The majority of differentially expressed downregulated genes were associated with ‘cell differentiation and morphology’ such as keratin (*KRT1*, *KRT4*, *KRT6B*, *KRT6C*, *KRT9*, *KRT23*, *KRT75*, *KRT77*, *KRT78*, and *KRT79*), filaggrin (*FLG*), envoplakin (*EVPL*), involucrin (*IVL*), periplakin (*PPL*), repetin (*RPTN*), transglutaminase (*TGM1* and *TGM3*), small proline-rich protein (*SPRR1A*, *SPRR1B*, *SPRR2A*, *SPPR2B*, *SPRR2D*, *SPRR2E*, *SPRR2G*, and *SPRR3*), calcium-binding protein (*S100A9*, *CALML5*), and lipase family member (*LIPN* and *LIPM*) (**Supplementary Fig. 3**).

**Figure 5.**
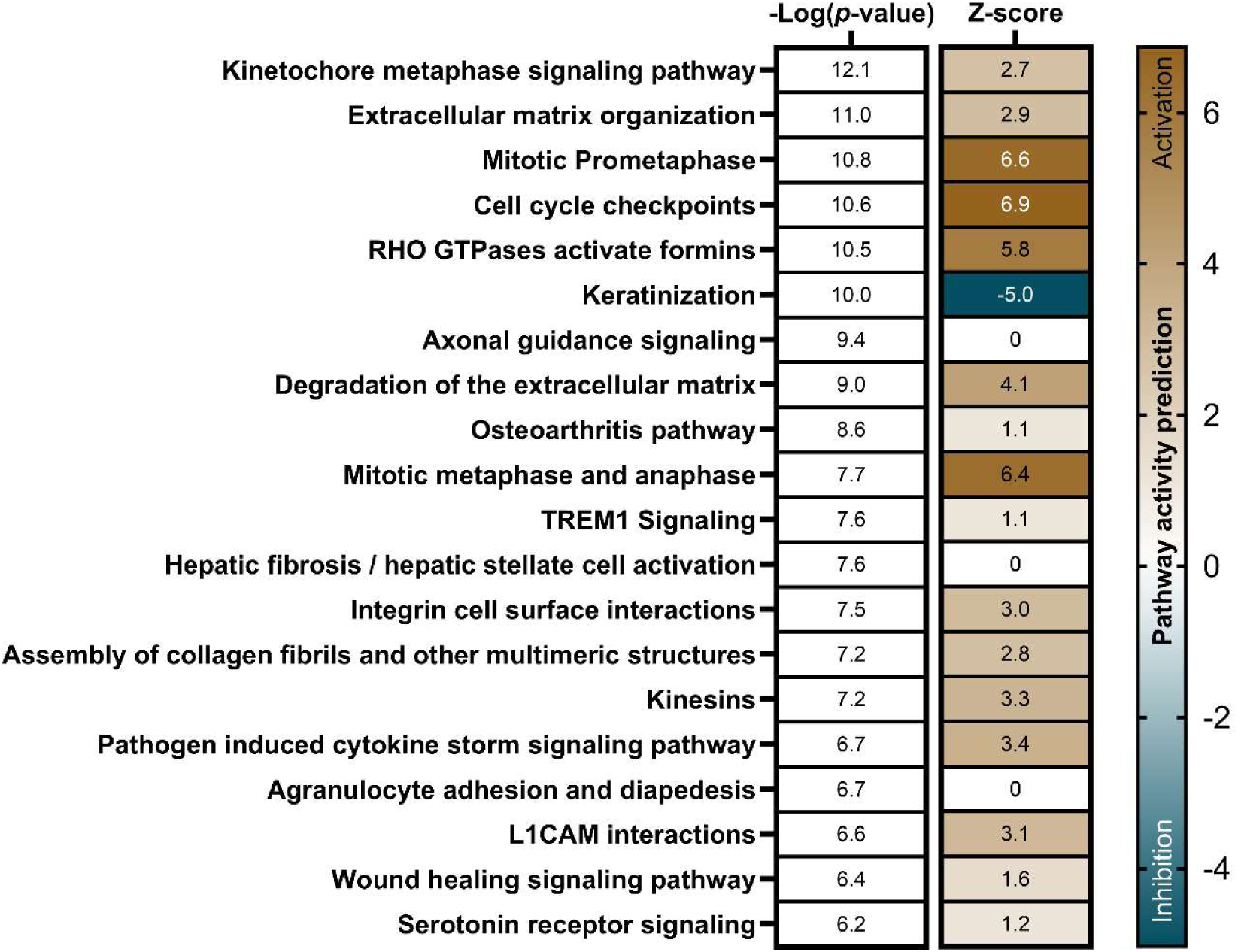
The top 20 significantly enriched biological pathways in keratinocytes co-cultured with *M. furfur* CBS 14141 versus monoculture. Cultures were performed on four independent occasions (N = 4), with a keratinocyte input of 1 × 10^5^ cells/well (10^4.5^ cells/cm^2^) and *Malassezia* of 1 × 10^6^ CFU/well (10^5.5^ CFU/cm^2^), and incubated for seven days. Biological pathways were ranked using the - log *p*-value of overlap. The -log *p*-value of overlap was derived from the right-tailed Fisher’s Exact test which determines the overlap between a subset of analysis-ready genes (± log_2_ fold-change of 2.5; FDR < 0.05) and a particular biological pathway with known gene functions. The Z-score represents the predicted biological pathway activity and was computed using a subset of analysis-ready genes and the known gene function in a specific biological pathway: higher (brown) suggests activation, lower (teal) suggests inhibition, and zero indicates no predicted activity. Pathway analysis was performed using Ingenuity Pathway Analysis software (QIAGEN). See **Supplementary Table 6** for a list of genes associated with specific pathways.

### Malassezia exposed to keratinocytes show perturbed metal, substrate, and protein-associated gene expression

Putative *M. furfur* metal-associated genes upregulated in co-cultures were ferric reductase (*GLX27_002799*), copper transporter complex subunit (*CTR4*), zinc-finger protein (*SUR1*), zip-like iron-zinc transporter (*GLX27_001560*), ferro-O_2_- oxidoreductase (*GLX27_004283*), and uroporphyrinogen decarboxylase (*HEM12*) (**Fig. 6**). Putative substrate transport-associated genes upregulated were the major facilitator superfamily general substrate transporters (*GLX27_003683*, *GLX27_003742,* and *GLX27_000029*) and sugar transporter (*GLX27_003485*). Most upregulated substrate transport genes were putative sugar transporters. Downregulated substrate transport genes were related to amino acid and pyrimidine transporters: general amino acid permease (*AGP2_1* and *AGP2_2*) and uracil permease (*GLX27_004590*) (**Fig. 6** and **Supplementary Fig. 4**). Putative protein metabolism-associated genes upregulated were prolyl oligopeptidase (*GLX27_000991*), carboxypeptidase D (*GLX27_002698*), glutamate decarboxylase (*GAD2*), 3-deoxy-7-phosphoheptulonate synthase (*GLX27_000589*), homocitrate synthase (*LYS4_2* and *LYS21_1*), acetolactate synthase (*ILV2*), asparagine synthase (*ASN1*), and aspartate-semialdehyde dehydrogenase (*HOM2*). Putative protein metabolism-associated genes downregulated were amidase (*GLX27_004500*), 3- hydroxyphenylacetate 6-hydroxylase (*GLX27_004595*), and primary-amine oxidase (*GLX27_003392*, *GLX27_002111*, and *GLX27_002109*).

**Figure 6.**
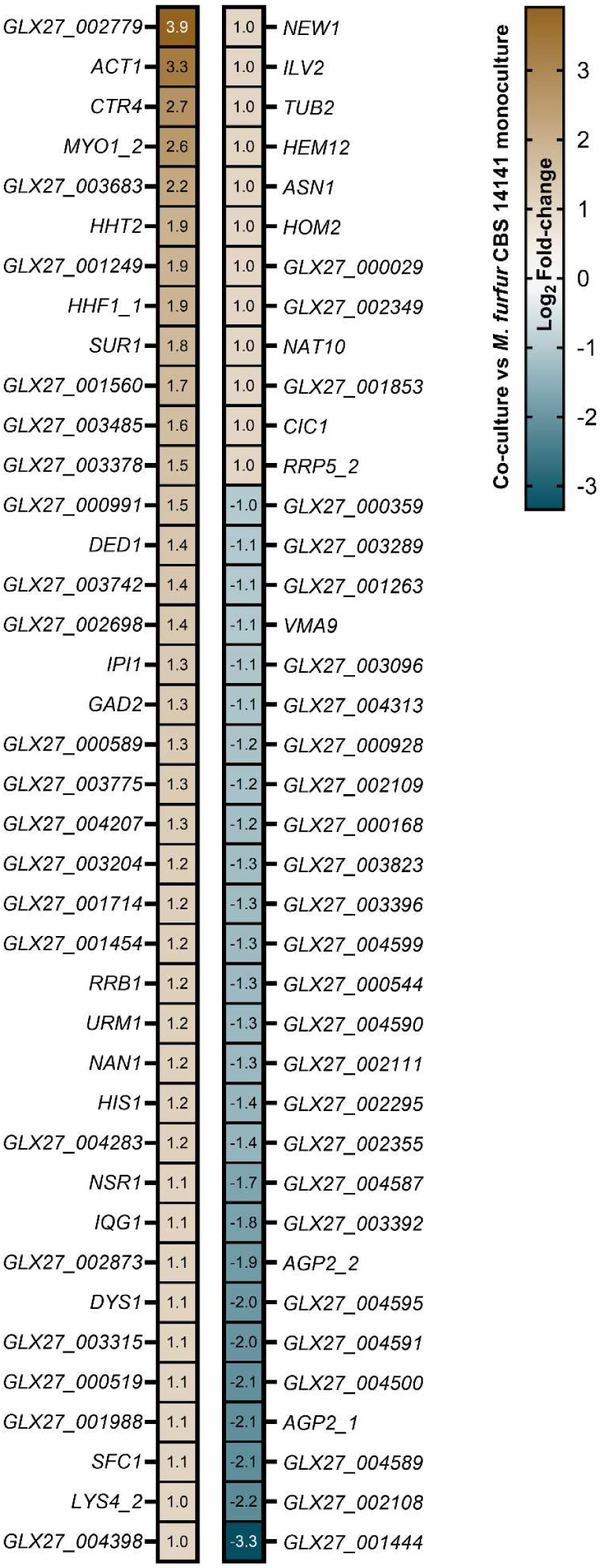
Gene expression profile of *M. furfur* CBS 14141 co-cultured with keratinocyte versus monoculture. Cultures were performed on four independent occasions (N = 4), with a keratinocyte input of 1 × 10^5^ cells/well (10^4.5^ cells/cm^2^) and *Malassezia* of 1 × 10^6^ CFU/well (10^5.5^ CFU/cm^2^), and incubated for seven days. Only analysis-ready genes (± log_2_ fold-change of 1; FDR < 0.05) are shown. Transcriptomics analysis was performed using *M. furfur* CBS 14141 complete genome with associated annotations (GenBank: GCA_009938135.1). Gene expression values are in log_2_ fold-change scale: upregulated genes in co-cultured *M. furfur* are in brown while downregulated expression is in teal.

## Discussion

While menopause impacts half of the world’s population, it remains poorly studied and understood, particularly its impact on skin homeostasis.^4–6,10,17–22^ The impact of menopause on the skin mycobiome, and vice versa, remains essentially unknown. In order to more objectively define menopausal stages and the associated changes in women’s skin and mycology we employed an objective method to stratify menopausal status based on blood hormone concentrations (**Fig. 1B** and **Supplementary Fig. 1**), ensuring robust classification for assessing associated skin physiological measures including TEWL, sebum, and pH, along with mycobiome composition.

Our findings revealed a reduction in TEWL (**Fig. 1G**) and sebum concentration (**Fig. 1H**) in post-menopausal compared to pre-menopausal women, consistent with previous studies.^50–53^ *Malassezia* was the most abundant fungal genus in both groups (**Fig. 2B**), with no significant differences in median relative abundance at the genus and species levels, likely due to the dominance of the fungi on human skin^54^. However, we observed variations in detection frequencies, most notably a ∼20% increase in *M. globosa* detection on post-menopausal women’s scalp (**Fig. 2D**). Broad-based cohort analyses using simple descriptive statistics may not fully capture the complex relationships between skin parameter changes and microbial abundance. Correlation analyses better encapsulate pair-wise relationships between two variables. Non-parametric spearman correlation identified *M. globosa* negatively correlating with cheek and scalp sebum concentration, while *M. restricta* positively correlating with cheek and scalp sebum (**Fig. 2E**). Given that post-menopausal women exhibit lower sebum concentrations, these correlations suggest that *M. globosa* is more abundant, while *M. restricta* is less abundant, on the cheeks and scalps of post-menopausal women.

In addition to changes in skin parameters, women experience a reduction in skin thickness as they progress toward menopause.^55^ The epidermis serves as the interface between the mycobiome and the deeper layers of the skin. The thinner epidermis in post-menopausal women is due to a lower keratinocyte population compared to pre-menopausal women.^55^ To investigate how changes in the epidermal cell population influence the interactions with *Malassezia*, we co-cultured human immortalized keratinocytes (N/TERT-1) with various skin-relevant *Malassezia* species at various cell population ratios. Keratinocytes tolerated a fungal load of 10⁵ CFU/well (10^4.5^ CFU/cm²), but cellular toxicity, indicated by drastically reduced keratinocyte viability, was observed at and beyond a fungal load of 10⁶ CFU/well (10^5.5^ CFU/cm²) (**Fig. 3A**). Among the *Malassezia* species tested, *M. globosa* exhibited the highest virulence toward keratinocytes (**Fig. 3B**), highlighting species-specific toxicities. Given that post-menopausal women produce less sebum and that lower sebum concentration correlates with higher *M. globosa* detection and relative abundance, the increased presence of *M. globosa* on post-menopausal skin may lead to deleterious effects.

Keratinocytes exposed to *Malassezia* exhibited an overproduction of chemokines, cytokines, and proteinases at the transcript and protein levels when co-cultured with a toxic *Malassezia* fungal load of 10⁶ CFU/well (10^5.5^ CFU/cm²) (**Fig. 4** and **Supplementary Fig. 3**). Additionally, *Malassezia* was detected intracellularly within co-cultured keratinocytes, in proximity to the nucleus, and demonstrated increasing *Malassezia* cell number from 24 hours to 7 days of culture, revealing intracellular replication and/or accumulation (**Fig. 3C**, **3D**, and **Supplementary Fig. 2**). The ability of *Malassezia* to infiltrate into keratinocytes may enable it to evade immune clearance while persistently recruiting inflammatory immune cells such as neutrophils and macrophages, leading to a chronically inflamed skin environment. Combined with its ability to cause keratinocyte cell death, an increased *Malassezia* to keratinocyte ratio in post-menopausal women may contribute to reduced epidermal keratinocyte population, and, coupled with elevated and sustained production of proteases^55,56^, lead to thinning skin.

Comparative transcriptomics of co-cultured keratinocytes revealed upregulation of genes associated with cell division pathways, while key genes involved in keratinocyte differentiation, such as *KRT*, *FLG*, *EVPL*, *IVL*, *TGM*, and *SPRR*, were markedly downregulated (**Fig. 5** and **Supplementary Fig. 3**). This suggests that keratinocytes co-cultured with *Malassezia* proliferate without undergoing proper differentiation. The *in vitro* model suggests that *Malassezia* may inhibit *in vivo* epidermal keratinocyte differentiation, disrupting the formation of the protective stratum corneum while promoting keratinocyte hyperproliferation, facilitating *Malassezia* intracellular invasion and replication within deeper epidermal layers. Notably, psoriasis^57,58^ and dandruff^33,59^ are characterized by undifferentiated, hyperproliferative keratinocytes in an inflammatory environment, resembling our *in vitro* keratinocyte/*Malassezia* co-culture model. The *in vitro* co-culture model system provides an avenue for further exploration into the host-microbe interaction and the development of targeted therapies to mitigate *Malassezia*-associated dermatological challenges in post-menopausal women.

For example, *NLRP3*, an inflammasome that triggers inflammatory programmed cell death, was overexpressed in co-cultured keratinocytes (**Supplementary Fig. 3**). Activation of NLRP3 can be determined by p17, the active secreted form of IL-1β,^49^ which was detected specifically in the supernatant of keratinocytes co-cultured with *Malassezia* (**Supplementary Fig. 5**). NLRP3 is implicated in various inflammatory skin conditions, including acne, dermatophytosis, psoriasis, and other immune-mediated skin diseases.^60,61^ Several NLRP3 inhibitors undergoing clinical evaluation^62^ may have potential as topical anti-inflammatory agents for *Malassezia*-induced skin inflammation, such as *Malassezia* folliculitis^63^. Similarly, *IL23*, a key cytokine in the IL23/IL17 axis involved in *Malassezia-*associated skin diseases like atopic dermatitis and psoriasis,^64–66^ was upregulated in co-cultured keratinocytes. While IL23 promotes inflammation, its activation of the IL17 pathway in γδ T cells has been shown to regulate *Malassezia* fungal load in a murine model.^65^ However, overproduction of IL23 and IL17 leads to inflammatory skin diseases, in which inhibitors of these cytokines serve as effective treatments.^67^ Therefore, while targeting NLRP3 and the IL23/17 axis presents a promising approach for *Malassezia*-associated inflammatory skin disease, formulations must carefully balance controlling the population of specific *Malassezia* species while preserving beneficial immune responses. The *Malassezia-*keratinocyte model reported herein serves as a platform to further investigate the interactions, enabling the development of anti-inflammatory pharma- and cosmeceuticals for post- menopausal women.

Understanding the *Malassezia* response to keratinocytes using transcriptomics remains challenging due to the lack of well-annotated genes, even in the model organism, *M. furfur*. Our analysis revealed that co-cultured *Malassezia* upregulated metal and substrate uptake genes, suggesting potential nutrient deprivation, while increased expression of protein metabolism-associated genes indicated an adaptive survival response (**Fig. 6**). However, these interpretations remain speculative. To gain deeper insights into *Malassezia* gene function and its interactions with keratinocytes, we are developing an *M. furfur* CBS 14141 transposon library, where isogenic mutants for all non-essential genes are created.

Lastly, *Malassezia* was detected within keratinocytes near its nucleus, suggesting potential localization within the endoplasmic reticulum, where lipids and proteins are synthesized (**Fig. 3D** and **Supplementary Fig. 2**). This raises the possibility that *Malassezia* hijacks keratinocytes to replicate within the endoplasmic reticulum while evading immune clearance. It is evident that *Malassezia* induces a proinflammatory response from keratinocytes (**Fig. 4**); infiltration within skin keratinocytes and immune evasion can contribute to chronic skin inflammation. Future research will investigate *Malassezia* intracellular localization in skin keratinocytes and identify specific subcellular sites to guide the development of intracellular antifungal treatments aimed at mitigating *Malassezia*-induced skin inflammation and promoting healthier skin in post-menopausal women.

## Conclusion

Women’s skin undergoes significant changes during menopause, including reduced thickness and sebum production, which then influence the mycobiome by altering the skin environment, nutrient availability, and consequently, host-microbe interactions. Lower sebum levels mean fewer lipids for lipid-dependent *Malassezia* species, leading to increased competition for nutrients and potentially shifting *Malassezia* from a commensal to a pathogenic state. Notably, detection of *M. globosa* and *M. furfur*, both associated with skin disease, was markedly higher in post-menopausal women. *In vitro*, keratinocytes tolerated *Malassezia* up to a specific fungal load, beyond which dose-dependent toxicity was observed. Among the tested species, *M. globosa* exhibited the highest virulence, causing significant keratinocyte cell death. Keratinocytes exposed to high fungal loads exhibited a transcriptome and cytokine secretome profile indicative of hyperproliferation and impaired differentiation in addition to inflammation, resembling patterns observed in psoriasis, seborrheic dermatitis, and other inflammatory skin conditions. Our *Malassezia*–keratinocyte co-culture model offers a valuable platform for dissecting host–microbe interactions, elucidating disease mechanisms, and supporting the development of targeted anti- fungal and anti-inflammatory therapies for *Malassezia*-associated skin disorders in post-menopausal women. Next steps include identifying *Malassezia* localization on post-menopausal women’s skin and unraveling how *Malassezia* invade keratinocytes to guide the development of targeted anti-fungal and anti-inflammatory therapies.

While this study primarily utilized metagenomics to assess *Malassezia* relative abundance, it does not provide absolute quantification in terms of cell population density or biomass. Future studies could incorporate quantitative polymerase chain reaction with species-specific standard curves (cell count versus internal transcribed spacer copy number) to estimate absolute abundance. Our findings identify key *Malassezia* species of interest that are differentially present on the cheek and scalp as women progress through different stages of menopause, warranting further investigation using more quantitative approaches.

## Materials and methods

### Participant recruitment

A total of 382 female participants aged 30 and above were recruited from the Health For Life In Singapore (HELIOS)^41^ cohort for this study, led by the Asian Skin Microbiome Program (ASMP). Participants were excluded from the study if they were pregnant, breastfeeding, had an ongoing illness, or had undergone a recent major surgery. Participants provided written informed consent before enrollment, and all procedures were conducted in accordance with the Declaration of Helsinki. The study was approved by the Nanyang Technological University (NTU) Institutional Review Board (IRB-2016-11-030) and the National Healthcare Group Domain Specific Review Board (D/09/021). Participants completed diagnostic questions and self-assessments regarding their perception of their skin health. A visual skin examination was conducted to assess psoriasis at the hairline. Additional metadata, including age, gender, race, and ethnicity, were collected. A total of 345 representative females were included in this study after excluding 37 females that could potentially confound our analyses (on prescription birth control [N = 8], had hysterectomy [N = 6], had hysterectomy and oophorectomy [N = 5], used hormone replacement therapy [N = 4], used Mirena IUD and stopped mensuration [N = 2], used unspecified IUD and mensuration stopped [N = 3], used unspecified medication and mensuration stopped [N = 4], had unspecified surgery and mensuration stopped [N = 3], had polycystic ovary syndrome [N = 1], and transsexual [N = 1]).

### Serum hormone quantification

Fasting blood samples were collected from participants in the morning by phlebotomists into 5 mL vacutainers (BD; Cat. #367955). Within minutes of collection, blood was transported to the laboratory, allowed to clot in an upright position at room temperature for 30 minutes before centrifugation at 2500g for 10 minutes at 4 °C. Vacuatiners were then loaded into an immunoassay system (SIEMENS; ADVIA Centaur XPT) to measure serum E2, FSH, and LH concentrations. In addition to daily system maintenance, calibrations were performed every 21, 14, and 28 days for E2, FSH, and LH, respectively.

### Skin parameter measurements

Participants acclimated to the sampling zone for at least 20 minutes under controlled conditions (ambient temperature: 20–22°C; relative humidity: 40–60%). TEWL was measured at the right temple using a VapoMeter (DEFLIN). Three consecutive measurements were taken at the same site, with a 10-minute interval between each to allow the skin to equilibrate with atmospheric air. Participants were instructed to refrain from consuming caffeinated beverages for at least three hours before sampling and to avoid applying topical products to the sampling site. Skin pH was measured at the left temple using a flat-bottom pH meter (DELFIN), with 20 µL of distilled water applied to the site before measurement. Three pH readings were taken at the same location. If participants had recently washed their face with tap water, synthetic detergents, or alkaline soaps, a delay of 3, 5, or 10 hours, respectively, was observed before sampling. For participants who had applied ointments, body lotions, or other topical products, pH measurement was delayed for 12 hours after the last application. Sebum levels were measured at the right cheekbone (over the zygomatic arch) using a sebum meter (DELFIN; SebumScale). A single measurement was recorded. Participants were advised to avoid applying topical products to the measurement area for at least 12 hours before sampling. For TEWL and pH measurements, triplicate readings were averaged for data analysis.

### Skin microbiome sampling

Circular adhesive disks (CLINICAL & DERM; Cat. #D100) with a surface area of 3.8 cm² were used to collect microbial biomass from the stratum corneum of the participants’ facial and scalp skin. Sampling was conducted on either the left or right side of the face and scalp. The adhesive disk was firmly pressed onto the skin surface using the thumb to ensure even contact, then removed and reapplied to the exact same location for a total of 25 reapplications. After sampling, the adhesive disk was carefully folded and placed into a 1.8 mL microfuge tube. Samples were stored at - 80°C until DNA extraction.

### Microbiome DNA extraction

Microbial genetic material from cells on the adhesive disks was extracted by first adding 500 µL of lysis buffer ALT (QIAGEN; Cat. #939011) and then transferred to a bead-beating tube (MP; Cat. #1169140-CF). Samples were subjected to bead-beating at 4 m/s for 30 seconds using an automated homogenizer (MP BIOMEDICALS; FastPrep-24). Beat-beating was performed twice per sample. The tubes were then centrifuged at maximum speed (16,000 RCF) for 5 minutes, resulting in 200 µL of supernatant, which was then transferred into 2 mL EZ1 sample tubes (QIAGEN) with 12 µL Proteinase K. The tubes were vortexed and incubated for 15 minutes at 56°C. DNA was extracted using magnetic beads (QIAGEN; Cat. #953034) with the help of an extraction instrument (QIAGEN; EZ1 Advance XL). Purified DNA was eluted in 50 µL elution buffer.

### Shotgun metagenomics

Library preparation for purified DNA was conducted using NEBNext Ultra II FS DNA Library Prep Kit for Illumina (NEW ENGLAND BIOLABS; Cat. #E7805) following the manufacturer’s protocol. DNA was enzymatically fragmented in a thermocycler at 37°C for 10 minutes, followed by 65°C for 30 minutes, generating DNA fragments ranging from 100 to 250 base-pairs. Illumina-compatible adaptors (NEW ENGLAND BIOLABS; Cat. #E6609) were ligated to the fragmented DNA, and adaptor-ligated fragments were purified using paramagnetic beads (BECKMAN COULTER; Cat. #A63882) at a 7:10 beads-to-sample ratio. Purified DNA fragments were then indexed with unique barcode sequences and subjected to 12 cycles of amplification per the manufacturer’s instructions. The quality of DNA fragments was assessed via electrophoresis (AGILENT; Cat. #5067-558), and samples passing quality control were normalized to ensure uniform DNA concentration and volume across all samples.

Processed DNA fragments underwent 150 base-pair paired-end sequencing on the Illumina HiSeq X platform, generating ∼12 million paired reads per sample. Sequence data were processed using CLC Genomics Workbench Premium (QIAGEN) version 23.0.5. Adaptor sequences were removed, and reads were quality-filtered using default parameters to trim low-quality base calls (CLC; Trim Reads 2.9). Human-derived reads were eliminated by mapping all reads to the human genome (GRCh38) (CLC; Taxonomic Profiling 2.36). The remaining non-human reads were aligned against a custom fungal genome database (**Supplementary Table 2**). To mitigate biases associated with varying library sizes, rarefaction analysis guided the subsampling of each sample to 10,000 paired reads. These subsampled were then mapped to the custom fungal genome database at a 99-seed length (99/150 base-pairs), the highest permitted by the CLC software. All subsequent taxonomic analyses were conducted using default settings within CLC Workbench’s microbial metagenomics template workflow.

### Keratinocyte and Malassezia culture

N/TERT-1 keratinocytes^68^ (passage 83) were seeded at a density of 600,000 cells per 175 cm² flask (NUNC; Cat. #178883) in 45 mL of keratinocyte serum-free medium (KSFM) (GIBCO; Cat. #17005042) supplemented with 25 µg/mL bovine pituitary extract and 0.2 ng/mL recombinant epidermal growth factor, herein referred to as KSFM. Cultures were incubated at 37°C with 5% CO₂ and humidification for three days. Upon reaching ∼80% confluency, spent media was discarded, and adherent cells were rinsed with 20 mL of phosphate-buffered saline (PBS) (GIBCO; Cat. #20012043). Cells were detached by adding 15 mL of recombinant trypsin (GIBCO; Cat. #12604-013) and incubating at 37°C for 8 minutes. Recombinant trypsin activity was neutralized with 15 mL of KSFM, and the 30 mL cell suspension was transferred to a 50 mL tube (NUNC; Cat. #339652). Cells were centrifuged at 300 RCF for 5 minutes, supernatant removed, and the pellet resuspended in 5 mL KSFM. Viability was assessed by mixing equal volumes of cell suspension and 0.4% trypan blue (GIBCO; Cat. #15250061), loading 10 µL onto a cell counting chamber slide (INVITROGEN; Cat. #C10283), and analyzing with an automated cell counter (INVITROGEN; Countess 3). Viable keratinocytes at specific densities were aliquoted into 12-well plates (NUNC; Cat. #150628) in 2 mL KSFM and incubated in a tissue culture incubator (ESCO; CelCulture CCL-170B-8) at 37°C for 24 hours to facilitate attachment to culture surfaces.

*Malassezia* species were cultured in 125 mL Erlenmeyer flasks (CORNING; Cat. #431143) containing 12 mL mDixon medium at 32°C with agitation (150 RPM) in a shake incubator (NEW BRUNSWICK SCIENTIFIC, Innova 44) for up to three days, depending on species growth profiles. After incubation, cells were transferred to a 50 mL tube, centrifuged at 1,250 RCF for 5 minutes, and the supernatant was discarded. The cell pellet was washed three times with 20 mL PBS. To achieve specific fungal seeding densities for mono- and co-cultures with keratinocytes, *Malassezia* suspensions were diluted in KSFM, ensuring the final PBS concentration remained below 1% of the total KSFM volume. For interaction studies, spent media from keratinocyte monocultures in 12-well plates (∼90 to 100% confluent) were replaced with 3 mL of fresh KSFM, while co-cultures received 3 mL of the diluted *Malassezia* suspension. Dilution accuracy was verified via CFU counts on mDixon agar (one CFU is initially derived from a single fungal cell). Mono- and co-cultures were incubated statically at 32°C with humidification and CO₂ (tissue culture incubator) for up to seven days.

### Keratinocyte and Malassezia viability assays

Post-culture supernatants (∼3 mL) were transferred into a new 12-well plate. Each culture well received 1 mL of PBS, followed by gentle agitation, after which the PBS was transferred to the 12-well plate containing the supernatants. The culture wells were then washed with an additional 1 mL of PBS, which was discarded. To detach keratinocytes, 1 mL of recombinant trypsin was added, and plates were incubated at 32°C for 10 minutes. Recombinant trypsin activity was neutralized by adding 1 mL of KSFM per well. The cell suspension was thoroughly resuspended, ensuring complete cell detachment. Keratinocyte viability was assessed using 0.4% trypan blue, as described previously. For *Malassezia* viability, post-culture supernatants were resuspended thoroughly, serially diluted in PBS, and spot-plated onto mDixon agar. Agar plates were incubated at 32°C until colonies were countable.

### Differential interference contrast and confocal microscopy

Post-culture supernatants were discarded, and wells were rinsed twice with 2 mL of PBS. Next, 1 mL of PBS was added to each well, and cultures were imaged using a differential interference contrast microscope (NIKON; ECLIPSE Ts2). Following imaging, PBS was replaced with 4% paraformaldehyde (GIBCO; Cat. #J61899-AK) and incubated at room temperature for 10 minutes to fix the cells. Wells were then rinsed twice with 2 mL of PBS. For staining, a mix was prepared in PBS, containing 1 µL of DAPI (SIGMA-ALDRICH; Cat. #MBD0015) and 1.5 µL of phalloidin (INVITROGEN; Cat. #A22287) per 1 mL of PBS. The wash PBS in each well was replaced with 1 mL of the staining mix and incubated at room temperature for 10 minutes. Wells were subsequently rinsed twice with 2 mL of PBS before a final addition of 2 mL of PBS for imaging with a confocal laser scanning microscope (OLYMPUS; FV3000RS).

### Multiplex microbead-based immunoassay

N/TERT-1 and *M. furfur* CRS 14141 mono- and co-culture supernatants (seven days post-culture) were filtered through a 0.22 µm pore-sized syringe filter (SARTORIUS; Cat. #S6534) and stored at -80°C. Cytokine analysis was based on the ProcartaPlex Human Immune Monitoring Panel, 65plex (INVITROGEN; Cat. # EPX650-16500-901). An aliquot of 50 µl of each sample was incubated with fluorescent-coded magnetic beads pre-coated with respective antibodies in a black 96-well clear-bottom plate overnight at 4°C. After incubation, plates were washed twice with wash buffer [PBS with 1% BSA (CAPRICORN SCIENTIFIC) and 0.05% Tween-20 (PROMEGA)].

Sample-antibody-bead complexes were incubated with biotinylated detection antibodies for one hour and washed twice with wash buffer. Subsequently, Streptavidin-PE was added and incubated for another 30 minutes. Plates were washed twice before sample-antibody-bead complexes were re-suspended in sheath fluid for acquisition on the FLEXMAP 3D (LUMINEX) using xPONENT 4.0 (LUMINEX) software. Data analysis was conducted on Bio-Plex Manager 6.1.1 (BIO-RAD). Standard curves were generated with a 5-parameter logistic algorithm, reporting values for mean florescence intensity and concentration data.

### RNA preservation, extraction, and quality control

Post-culture supernatant containing planktonic cells (∼3 mL) was combined with 9 mL of cold TRIzol reagent (AMBION; Cat. #15596018). An additional 1 mL of TRIzol reagent was used to lyse adherent keratinocyte cells within the 12-well plate cultures, and this suspension was pooled with the post-culture supernatant. Samples were stored at -80°C and thawed on ice before RNA extraction. For complete cell lysis via bead-beating, 0.5 mm zirconia/silica beads (BIOSPEC; Cat. #11079105z) were added up to the 0.5 mL mark in polypropylene tubes (LABCON; Cat. #3661-870-000-9). Each tube received 1 mL of TRIzol sample, and cells were lysed at 6.5 m/s for one minute using an automated homogenizer (MP BIOMEDICALS; FastPrep-24), followed by incubation on ice for five minutes. This bead-beating process was repeated five times. Lysates were then mixed with an equal volume of 100% molecular-grade ethanol (FISHER SCIENTIFIC; Cat. #BP2818500). RNA was extracted using the Direct-zol RNA Miniprep Kit (ZYMO; Cat. #R2052) following the manufacturer’s instructions and eluted with 25 µL of nuclease-free water (BIOBASIC; Cat. #WW1002). To minimize DNA contamination, samples were treated with the Turbo DNA-free Kit (AMBION; Cat. #AM1907), followed by purification with the RNA Clean & Concentrator-5 Kit (ZYMO; Cat. #R1014) and elution in 15 µL of nuclease-free water. RNA purity was assessed using a spectrometer (THERMO FISHER; NanoDrop One), while RNA and DNA concentrations were quantified using Qubit RNA BR (INVITROGEN; Cat. #Q10211) and Qubit 1X dsDNA HS (INVITROGEN; Cat. #Q33231), respectively, in Qubit assay tubes (INVITROGEN; Cat. #Q32856) and measured with a Qubit 4 fluorometer (INVITROGEN). RNA integrity was determined using RNA ScreenTape (AGILENT TECHNOLOGIES; Cat. #5067-5576) with the corresponding sample buffer (AGILENT TECHNOLOGIES; Cat. #5067-5577) on the TapeStation 4150 (AGILENT TECHNOLOGIES). The median quality values for all RNA samples were as follows: A260/280 = 2.14, A260/230 = 2.23, DNA contamination = 13.4%, and RNA integrity number = 8.

### RNA-sequencing and comparative transcriptomics

A total of 2 μg of RNA per sample was used as input material for RNA sequencing library preparation. Ribosomal RNA (rRNA) was first removed using the Ribo-Zero Plus rRNA Depletion Kit (ILLUMINA; Cat. #20037135), and the rRNA-depleted RNA was recovered through ethanol precipitation. Using the rRNA-depleted RNA, sequencing libraries were constructed with the NEBNext Ultra II Directional RNA Library Prep Kit for Illumina (NEW ENGLAND BIOLABS; Cat. #7760), following the manufacturer’s protocol. Briefly, RNA fragmentation was performed using divalent cations under elevated temperatures in the NEBNext First Strand Synthesis Reaction Buffer (5X). First-strand cDNA synthesis was carried out using random hexamer primers and M-MuLV Reverse Transcriptase (RNase H-). Second-strand cDNA synthesis followed, utilizing DNA Polymerase I and RNase H, with dNTPs containing dUTP instead of dTTP to facilitate strand specificity. The remaining overhangs were converted into blunt ends through exonuclease and polymerase activities. After adenylation of 3’ ends of DNA fragments, NEBNext Adaptor with hairpin loop structure were ligated to prepare for hybridization. To preferentially select cDNA fragments of 250–300 bp in length, the library fragments were purified using the AMPure XP system (BECKMAN COULTER; Cat. #A63880). The selected, adaptor-ligated cDNA was then treated with 3 μL of USER II Enzyme (NEW ENGLAND BIOLABS; Cat. #M5508) at 37°C for 15 minutes followed by 95°C for 5 minutes. Subsequently, PCR amplification was performed using Phusion High-Fidelity DNA Polymerase (THERMO FISHER; Cat. #F530S) with Universal PCR primers and Index (X) Primer. The final PCR products were purified using the AMPure XP system, and library quality was assessed using the Agilent Bioanalyzer 2100 system.

Processed cDNA fragments underwent 150 base-pair paired-end sequencing on the NovaSeq X Plus platform (ILLUMINA), generating approximately 20 million paired reads per sample. Sequencing reads were processed using CLC Genomics Workbench Premium (QIAGEN, version 23.0.5). Adapter sequences were removed, and reads were quality-filtered using default parameters to trim low-quality base calls. Quality-filtered reads were analyzed using the built-in RNA-Seq Analysis 2.7 following default settings for the RNA-Seq and Differential Gene Expression Analysis workflow. To identify differentially expressed genes in N/TERT-1 keratinocytes, sequencing reads were mapped to the human genome (GRCh38). For *M. furfur* CBS 14141, sequencing reads were mapped to the reference genome (GenBank: GCA_009938135.1), utilizing either publicly available annotations or in-house annotations. For pathway analysis, differentially expressed genes in N/TERT-1 with a Log_2_ fold-change of ±2 and a FDR p-value < 0.05 were analyzed using Ingenuity Pathway Analysis software (QIAGEN, version 24.0.1).

### *Malassezia* gene annotation search

Using published genome assembly and gene annotations (GCA_009938135.1), we determined differentially expressed *M. furfur* CBS 14141 genes (**Fig. 6**). Additionally, an in-house gene annotation for the same genome identified a subset of differentially expressed genes unique to co-cultured *M. furfur* CBS 14141 (**Supplementary Fig. 4**). However, multiple differentially expressed genes were annotated as hypothetical proteins. Therefore, we translated the gene nucleotide sequences to peptide sequences and queried the NCBI’s nr database to determine the likely identity of some of the differentially expressed genes. Herein are the gene name/locus of the differentially expressed genes and their associated homologous hit information from NCBI nr database: GLX27_002799 (ferric reductase; *M. pachydermatis*; coverage [c]: 99%; identity [i]: 37%), CTR4 (copper transporter complex subunit; *M. furfur*; c: 99%; i: 100%), SUR1 (zinc-finger protein ZAP1; *M. furfur*; c: 96%; i:98), GLX27_001560 (zip-like iron-zinc transporter; *M. pachydermatis*; c: 98%; i: 64%), GLX27_004283 (ferro-O2-oxidoreductase FET3; *M. furfur*; c: 90%; i: 100%), HEM12 (uroporphyrinogen decarboxylase; *M. brasiliensis*; c: 99%; i: 95%), GLX27_003683 (major facilitator superfamily [MFS] general substrate transporter; *Moesziomyces antarcticus*; c: 84%, i: 36.9%), GLX27_003485 (sugar transporter STL1; *M. restricta*; c: 93%; i: 67%), GLX27_003742 (MFS general substrate transporter; *M. pachydermatis*; c: 98%; i: 60%), GLX27_000029 (MFS sugar transporter; *M. pachydermatis*; c: 97%; i: 65%), AGP2_1 (general amino acid permease; *M. furfur*; c: 91%; i: 98%), AGP2_2 (general amino acid permease; *M. furfur*; c: 98%; i: 100%), GLX27_004590 (uracil permease; *M. pachydermatis*; c: 99%; i: 70%), GLX27_000991 (prolyl oligopeptidase; *M. restricta*; c: 85%; i: 36%), GLX27_002698 (carboxypeptidase D; *M. furfur*; c: 99%; i: 100%), GAD2 (glutamate decarboxylase; *M. furfur*; c: 99%; i: 100%), GLX27_000589 (3-deoxy-7-phosphoheptulonate synthase; *M. brasiliensis*; c: 98%; i: 96%), LYS4_2 (homocitrate synthase; *M. furfur*; c: 99%; i: 98%), ILV2 (acetolactate synthase; *M. furfur*; c: 99%; i: 100%), ASN1 (asparagine synthase; *M. brasiliensis*; c: 97%; i: 97%), HOM2 (aspartate-semialdehyde dehydrogenase; *M. brasiliensis*; c: 99%; i: 98%), LYS21_1 (homocitrate synthase; *M. furfur*; c: 87%; i: 100%), GLX27_004500 (amidase; *M. furfur*; c: 97%; i: 93%), GLX27_004595 (3-hydroxyphenylacetate 6-hydroxylase; *M. furfur*, c: 96%; i: 100%), GLX27_003392 (primary-amine oxidase; *M. psittaci*; c: 98%; i: 81%), GLX27_002111 (primary-amine oxidase; *M. japonica*; c: 99%; i: 78%), and GLX27_002109 (primary- amine oxidase; *M. brasiliensis*; c: 87%; i: 96%).

### Statistical analyses

The specific statistical analyses performed are detailed in the figure descriptions. Briefly, statistical tests were conducted using GraphPad Prism (version 10.0.3 [403]). The choice between parametric and non-parametric tests was determined based on data normality, assessed using a QQ plot. If the data were normally distributed, parametric tests were applied; otherwise, log transformation was performed to achieve normality. If normality could not be attained, non-parametric tests were used. For comparative transcriptomics, statistical analyses were conducted using the built-in workflows within CLC Genomics Workbench Premium (QIAGEN, version 23.0.5) and Ingenuity Pathway Analysis software (QIAGEN, version 24.0.1).

## Acknowledgments

The authors thank the Agency for Science, Technology and Research (A*STAR)’s Singapore Immunology Network (SIgN) Multiplex Analysis of Proteins (MAP) platform for enabling the Luminex work for this publication. A*STAR’s SIgN MAP platform is supported by research grants including the Biomedical Research Council (BMRC) grant and the National Research Foundation (NRF), Immunomonitoring Service Platform (Ref: ISP: NRF2017_SISFP09) grant. The authors also thank Dr. Lim Ying Shiang for his advice and in interpreting the cytokine profiles and Mr. Lam Yuen In Hilbert for assisting in the in-house *Malassezia* genome annotation.

HELIOS is supported by Singapore Ministry of Health’s (MOH) National Medical Research Council (NMRC) under its OF-LCG funding scheme (MOH-000271-00), Singapore Translational Research (StaR) funding scheme (NMRC/StaR/0028/2017), the National Research Foundation, Singapore through the Singapore MOH NMRC and the Precision Health Research, Singapore (PRECISE) under the National Precision Medicine programme (NMRC/PRECISE/2020), National Cohorts Office (P2022-02- 03) and intramural funding from Nanyang Technological University, Lee Kong Chian School of Medicine and the National Healthcare Group. The HELIOS team is also supported by a team of outstanding operational and administrative staff. This research was also supported by the Singapore National Research Foundation under its Translational and Clinical Research (TCR) Flagship Programme and administered by the Singapore Ministry of Health’s National Medical Research Council (NMRC), Singapore - NMRC/TCR/004-NUS/2008; NMRC/TCR/012-NUHS/2014. Additional funding was provided by the Singapore Institute for Clinical Sciences, Agency for Science Technology and Research (A*STAR), Singapore. This study was also supported by Agency for Science, Technology and Research (A*STAR) BMRC EDB IAF-PP grant (H17/01/a0/004) (TD); Skin Research Institute of Singapore, IAF-PP (HBMS) grant; Asian Skin Microbiome Program IAF-PP grants (H18/01/a0/016) (TD) and (H22/J1/a0/040); A*STAR BMRC Central Research Fund Translational Research (CRF-ATR) Award. JCC is supported by the Singapore Translational Research (StaR) funding scheme (NMRC/StaR/0028/2017) from 2017 - 2022. PR is funded by the National Research Foundation Fellowship, Singapore (NRF-NRFF11-2019-0006), and Nanyang Assistant Professorship (NAP). JC receives support from NIHR Newcastle Biomedical Research Centre.

The HELIOS Study Team is comprised of Prof Paul Eillott, Prof Eng Sing Lee, Prof Jimmy Lee, Prof Joanne Ngeow, Prof Sabrina Wong, Prof Elio Riboli, Dr Tricia Chang, Prof Rinkoo Dalan, Dr Wai Kee Kok, Dr Benjamin Lam, Dr Kelvin Li, Prof Tock Han Lim, Dr Pritesh Jain, Dr Hong Kiat Ng, Dr Theresia Mina, Dr Nilanjana Sadhu, Dr Akash Bahai, Dr Dorrain Low, Dr Xiaoyan Wang, Dr Harinakshi Sanikini, Dr Darwin Tay, Dr Terry Tong, Dr Kostas Tsilidis, Prof Wansaicheong Khin-lin, Gervais, in addition to Prof John Chambers, Prof Marie Loh, and Prof Yik Weng Yew.

## Author contributions

BD and TLD designed the *in vitro* studies, while the HELIOS Study Team, JC, JCC, YWY, ML, NN, and TLD designed the human studies. BD, JKR, AR, PR, ANMN, and NC conducted the experiments. BD, JKR, AR, PR, CL, NC, and TLD analyzed the data. BD and TLD developed the results and wrote the manuscript. All authors edited, read, and approved the final manuscript.

## Data availability

We will upload RNA seq raw sequencing files, but not skin shotgun metagenomic reads, because these genomic reads will be uploaded through an upcoming publication by Genome Institute of Singapore.

## Ethics declarations

All other authors declare no competing interests.

---

**Supplementary Figure 1.**
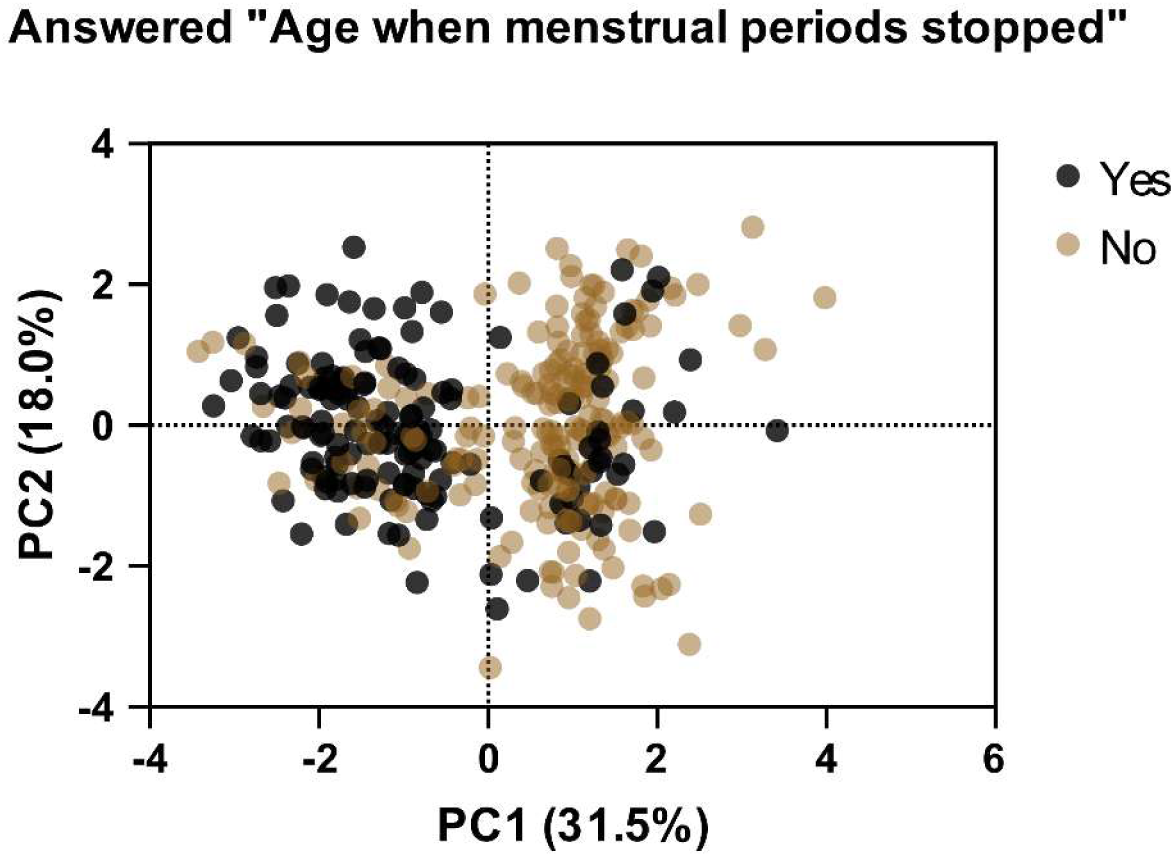
Principal component analysis of female serum hormone and skin physiology measures with a color overlay based on subjects’ responses to the menopause questionnaire. Each dot represents an individual subject. Black dots: ‘Yes’; Brown dots: ‘No’.

**Supplementary Figure 2.**
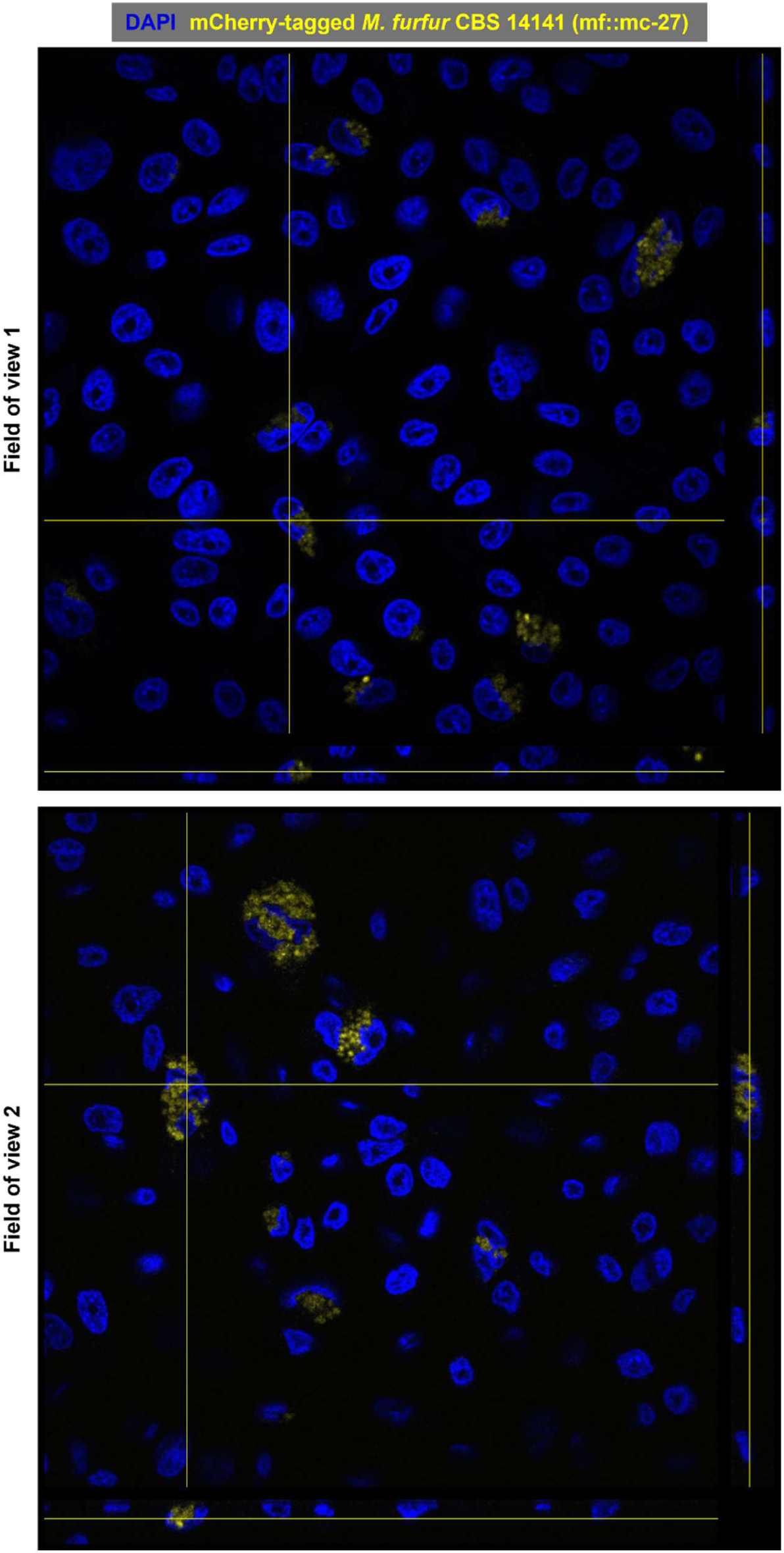
Keratinocytes (N/TERT-1) co-cultured with live mCherry-tagged *M. furfur* CBS 14141 (mf::mc-27) for seven days. Samples were fixed with 4% paraformaldehyde and stained with DAPI (binds to adenine-thymine-rich DNA regions; pseudocolored blue). mf::mc-27 was pseudocolored yellow. The vertical axis on the right-most subpanel shows the sagittal plane of the reconstructed Z-stack target cross-section, while the horizontal axis on the bottom-most subpanel shows the frontal plane. Confocal microscopy images are representative of three independent repeats and were acquired at 20× magnification.

**Supplementary Figure 3.**
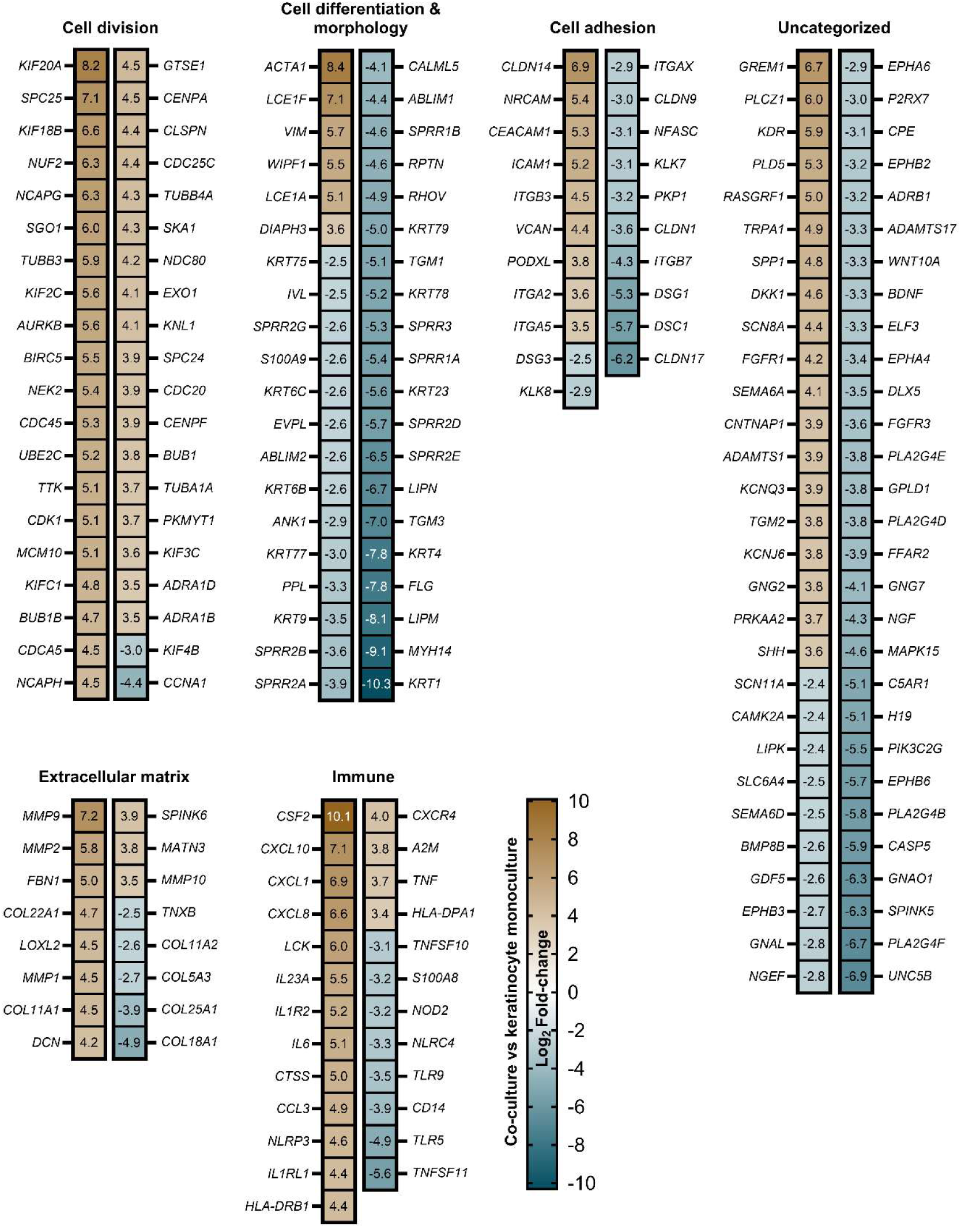
The 100 most significantly upregulated and downregulated genes in keratinocytes co-cultured with *M. furfur* CBS 14141. These genes were identified based on the ‘Top 20 significantly perturbed cellular functions/pathways’ presented in Figure 5. Differential expression was determined using a threshold of at least ± 2.5 log₂ fold-change with an FDR < 0.05.

**Supplementary Figure 4.**
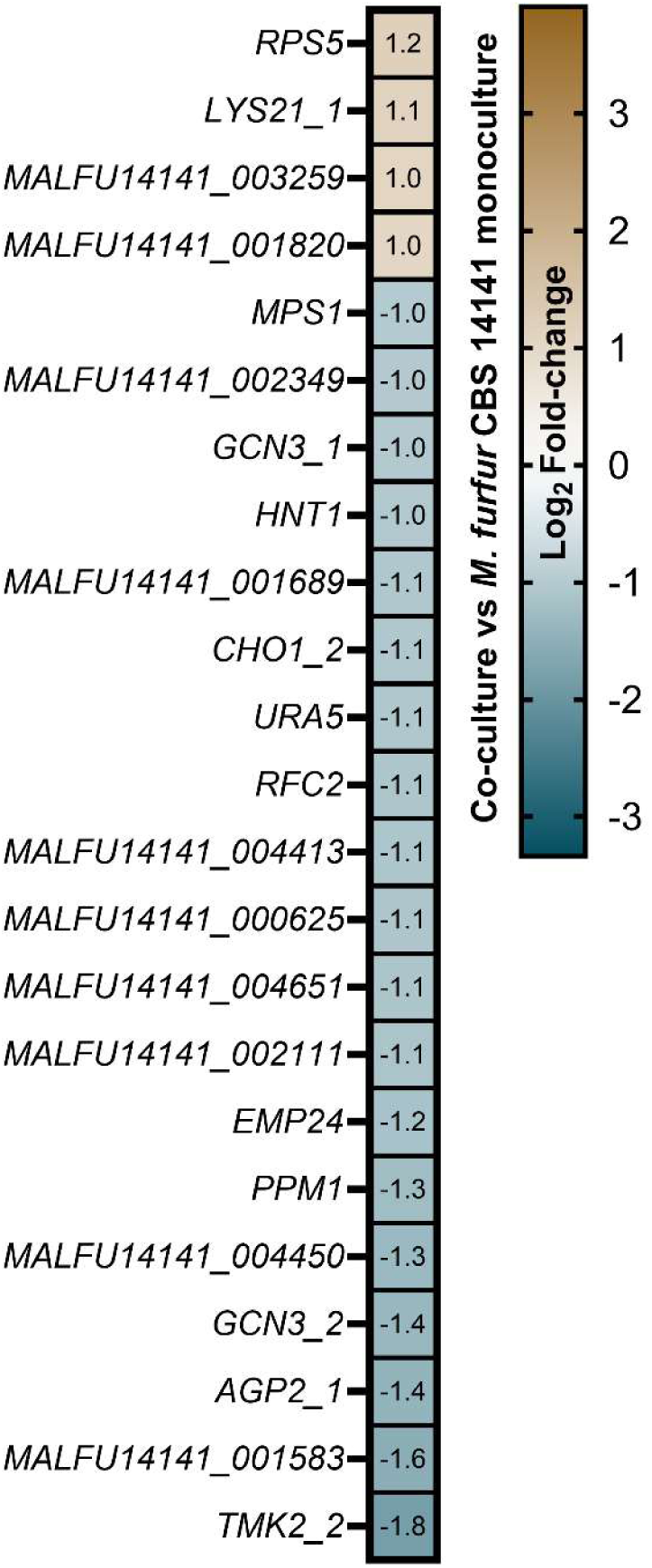
Additional differentially expressed genes in *M. furfur* CBS 14141 co-cultured with keratinocytes versus monoculture using an in-house annotation with GCA_009938135.1. Cultures were conducted in four independent experiments (N = 4) with a keratinocyte input of 1 × 10^5^ cells/well (10^4.5^ cells/cm2) and *Malassezia* of 1 × 10^6^ CFU/well (10^5.5^ CFU/cm2), incubated for seven days. Only analysis-ready genes (± log_2_ fold-change of 1; FDR < 0.05) and unique gene annotations absent from the published GCA_009938135.1 complete genome are shown. Gene expression values are in log_2_ fold-change scale: upregulated genes in co-cultured *M. furfur* are in brown while downregulated expression is in teal.

**Supplementary Figure 5.**
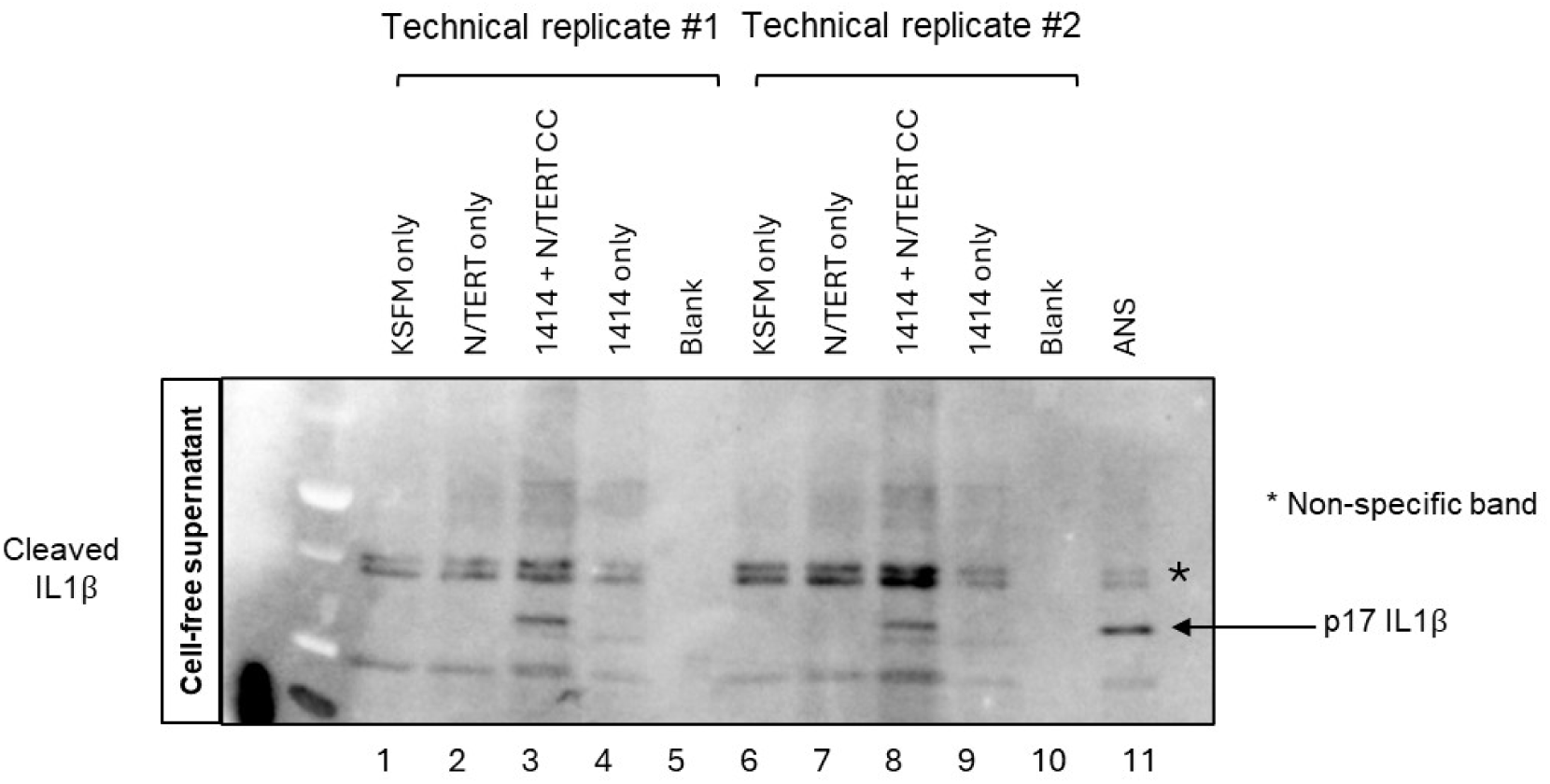
*M. furfur* induces p17 secretion by N/TERT-1. Co-culture supernatants were derived from seven-day cultures of *M. furfur* CBS 14141 and N/TERT-1. Supernatants were syringe-filtered through 0.2 µm pore-sized hydrophilic filters, and the cell-free supernatants were stored at -80 °C before analysis. Before analyses, frozen supernatants were thawed on wet ice, concentrated 10X (500 µL to 50 µL), and probed for p17 IL-1β (western blot). As a positive control for p17 secretion, N/TERT-1 was treated overnight with 1 µM anisomycin (ANS), and the supernatant was probed for p17.

